## Supplementary for "Sex-specific transcriptomic responses to changes in the nutritional environment"

**Table S1:** Compositions of essential and non-essential amino acid stock solutions.

| Amino acid stock solution |  | **(g/200 ml)** |
| --- | --- | --- |
| **Essential amino acid** | | |
| F (L-phenylalanine) |  | **3.03** |
| H (L-histidine |  | **2.24** |
| K (L-lysine) |  | **5.74** |
| M (L-methionine) |  | **1.12** |
| R (L-arginine) |  | **4.70** |
| T (L-threonine) |  | **4.28** |
| V (L-valine) |  | **4.42** |
| W (L-tryptophan) |  | **1.45** |
| **Non-essential amino acid** | | |
| A (L-alanine) |  | **5.25** |
| D (L-aspartate) |  | **2.78** |
| G (glycine) |  | **3.58** |
| N (L-asparagine) |  | **2.78** |
| P (L-proline) |  | **1.86** |
| Q (L-glutamine) |  | **6.02** |
| S (L-serine) |  | **2.51** |

**Table S2:** Recipe for 200ml of protein solution.

|  |  |  | **Total volume 200ml** |
| --- | --- | --- | --- |
|  | L-ile | Powder | 348mg |
|  | L-leu | Powder | 492mg |
|  | L-tyr | Powder | 252mg |
|  | cholesterol | 20mg/ml in EtOH | 3ml |
|  | CaCl2 | 1000x | 200ul |
|  | MgSO4 | 1000x | 200ul |
|  | CuSO4 | 1000x | 200ul |
|  | FeSO4 | 1000x | 200ul |
|  | MnCl2 | 1000x | 200ul |
|  | ZnSO4 | 1000x | 200ul |
|  | H_2_O |  | Up to 50ml |
|  | Total volume before autoclaving | | 50 ml |
|  | buffer | 10x acetate buffer base | 20ml |
|  | nucl/lipid soln | 125x stock | 1.6ml |
|  | Yaa solutions | essential amino acid stock solution (EAA) | 18.154ml |
|  |  | non-essential amino acid stock solution (NEAA) | 18.154ml |
|  |  | Na glutamate solution (100mg/ml) | 5.464ml |
|  |  | Cys solution (50mg/ml) | 1.584ml |
|  | Vitamin stock | 47.6x stock | 4.2ml |
|  | folic acid stock | 1000x stock | 200ul |
|  | Propionic acid |  | 1.2ml |
|  | Nipagin | 100 g/l stock in 95% EtOH | 3ml |
|  |  | Make to total volume of 200ml with H_2_O |  |

**Table S3:** Recipe for 200ml of carbohydrate solution.

|  |  |  | **Total volume 200ml** |
| --- | --- | --- | --- |
|  | sucrose | To match protein 1:1 | 6.5g |
|  | cholesterol | 20mg/ml in EtOH | 3ml |
|  | CaCl2 | 1000x | 200ul |
|  | MgSO4 | 1000x | 200ul |
|  | CuSO4 | 1000x | 200ul |
|  | FeSO4 | 1000x | 200ul |
|  | MnCl2 | 1000x | 200ul |
|  | ZnSO4 | 1000x | 200ul |
|  | H_2_O |  | Up to 50ml |
|  | Total volume before autoclaving | | 50ml |
|  | buffer | 10x acetate buffer base | 20ml |
|  | nucl/lipid soln | 125x stock | 1.6ml |
|  | Vitamin stock | 47.6x stock | 4.2ml |
|  | folic acid stock | 1000x stock | 200ul |
|  | Propionic acid |  | 1.2ml |
|  | Nipagin | 100 g/l stock in 95% EtOH | 3ml |
|  |  | Make to total volume of 200ml with H_2_O |  |

**Table S4:** Statistical analysis for the Experiment 3 (rapamycin treatment).

|  | Df | Sum Sq | Mean Sq | F value | Pr(>F) |
| --- | --- | --- | --- | --- | --- |
| sex | 1 | 0 | 0 | 0 | 1 |
| rapamycin | 3 | 40.259 | 13.4197 | 17.5106 | < 0.001 |
| diet | 1 | 9.026 | 9.0256 | 11.7769 | 0.0006704 |
| sex × rapamycin | 3 | 20.782 | 6.9274 | 9.0392 | < 0.001 |
| sex × diet | 1 | 8.654 | 8.6538 | 11.2918 | 0.0008631 |
| rapamycin × diet | 3 | 12.724 | 4.2412 | 5.5341 | 0.0010085 |
| sex × rapamycin × diet | 3 | 5.492 | 1.8308 | 2.3889 | 0.0686147 |
| Residuals | 355 | 272.063 | 0.7664 |  |  |

**Table S5:** Overlap between gene response classifications based on diet (our study) and IIS/TOR manipulation (Graze *et al.* [30]).

|  | D | D×S | D+D×S | None |
| --- | --- | --- | --- | --- |
| InR | 160  (-5%) | 1  (-89%) | 27  (+16%) | 2363  (-1%) |
| InR×S | 29  (+106%) | 1  (+29%) | 7  (+258%) | 177  (-10%) |
| InR+InR×S | 221  (+97%) | 15  (+143%) | 28  (+79%) | 1443  (-8%) |
| None | 137  (-46%) | 13  (-6%) | 14  (-60%) | 3674  (+4%) |

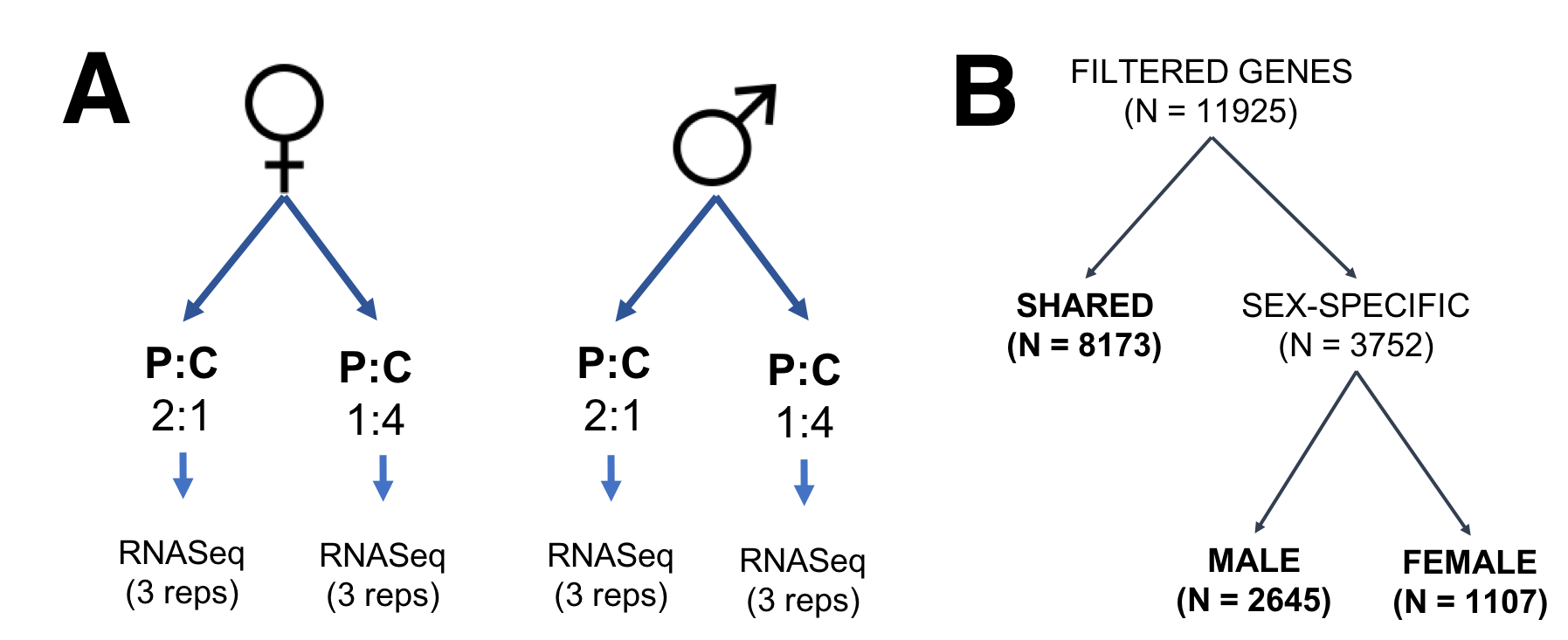

**Figure S1: (A)** Set-up for the transcriptomic experiment. **(B)** Experimental design for data analyses. Gene sets were split into three categories: those expressed in both sexes (shared) and those expressed in one sex only (sex-specific), further separated in those that are male-specific and female-specific in expression.

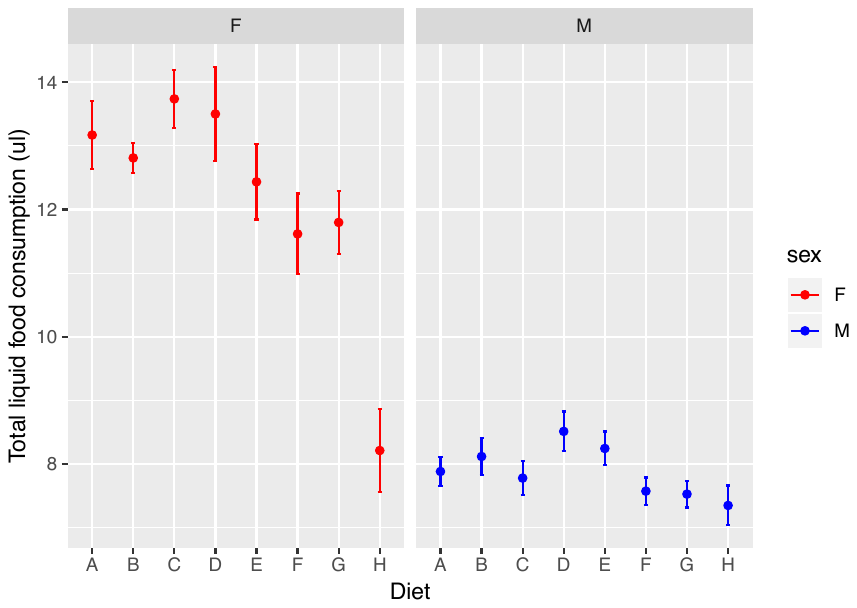

**Figure S2:** Total liquid diet consumption. Data for females is shown in red on the left and data for males in blue on the right. Diet composition ranges from A = high protein (P:C = 4:1) to H = high carbohydrate (P:C = 1:32).

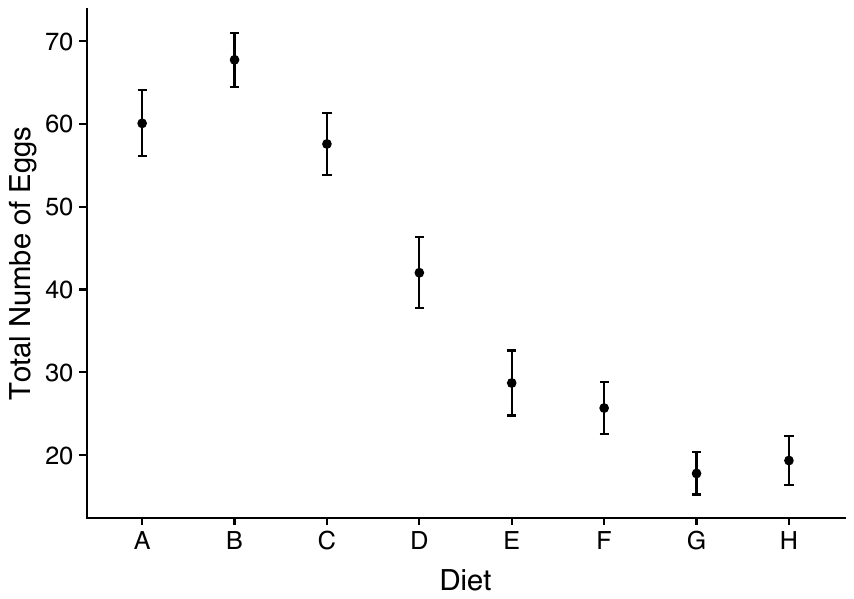

Female Fecundity

**Figure S3:** Female fecundity (number of eggs laid) across dietary treatments. Diet composition ranges from A = high protein (P:C = 4:1) to H = high carbohydrate (P:C = 1:32).

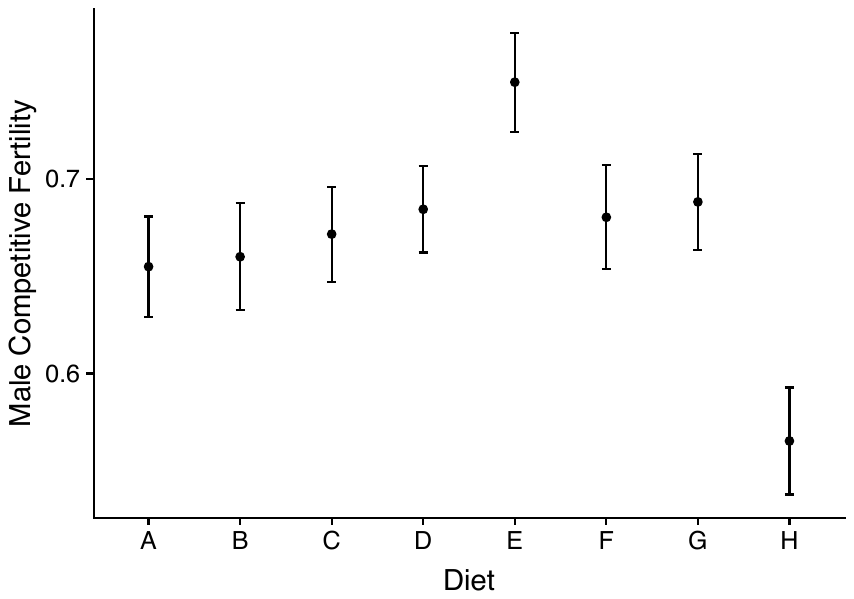

**Figure S4:** Male competitive fertility across dietary treatments. Diet composition ranges from A = high protein (P:C = 4:1) to H = high carbohydrate (P:C = 1:32)

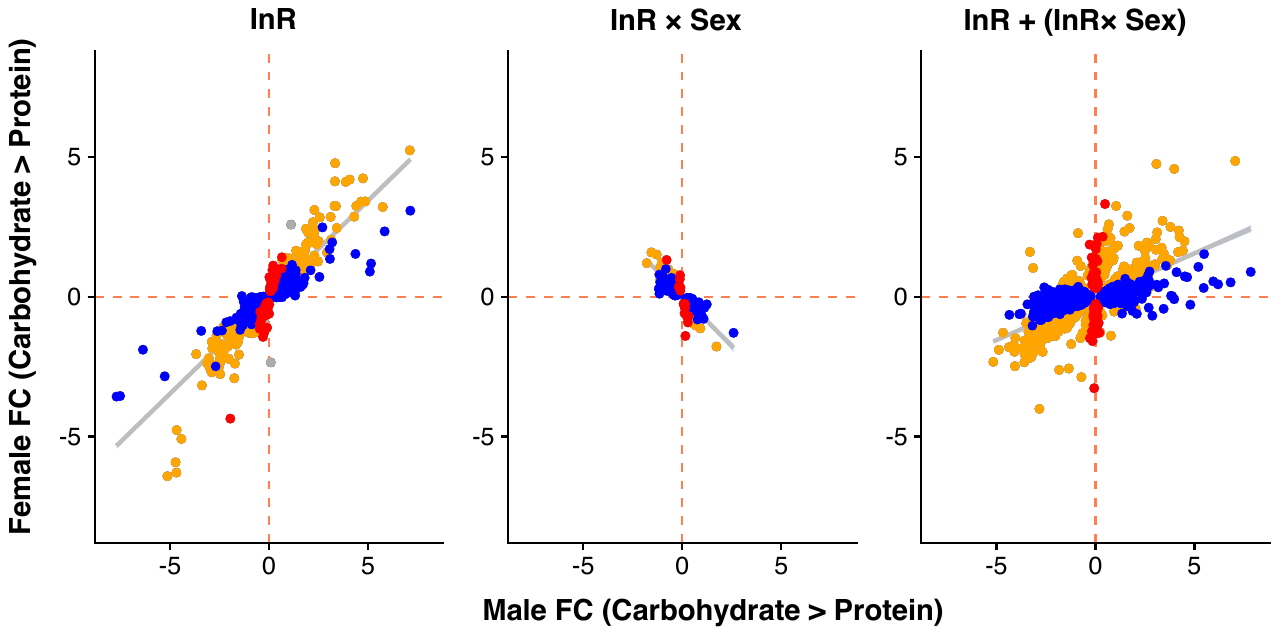

**Figure S5:** Male and female expression responses (log_2_-fold change) for genes classified as showing only a diet effect (Diet), only a InR-by-sex interaction (InR×Sex) or both (InR + InR×Sex) in the re-analysis of the Graze *et al.* [30] dataset. Expression changes are measured from carbohydrate- to protein-rich diet. Colours represent genes with significant differential expression (at 5% FDR) only in females (red), only in males (blue), in both sexes (yellow) or in neither sex (grey).

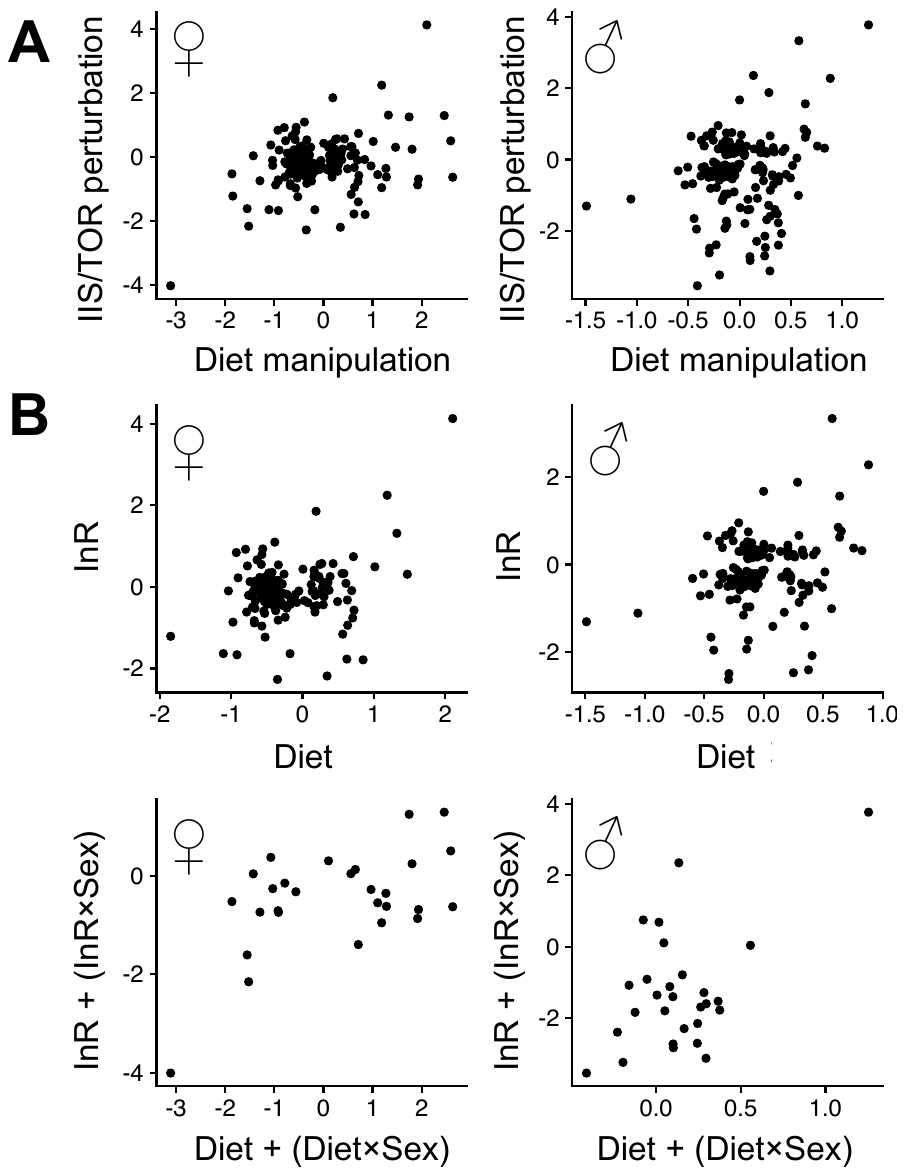

**Figure S6:** Female (left) and male (right) expression responses (log_2_-fold change) in response to IIS/TOR perturbation (Graze *et al.* [30] dataset) and diet manipulation. Panel (A) shows genes that with significant responses in both experiments (at 5% FDR, N=489). IIS/TOR and diet fold changes are significantly positively correlated in both sexes (Pearson's moment correlations, females: r = 0.32, p < 0.001, males: r = 0.20, p = 0.006). Panel (B) shows genes that fall in coinciding response classes (top: 'InR' and 'D', N=160; bottom: 'InR+ InR×Sex' and 'D+D×Sex', N=28, see Table S5). Fold changes are significantly positively correlated in all cases ('InR' and 'D' – females: r = 0.28, p < 0.001, males: r = 0.26, p < 0.001; 'InR+ InR×Sex' and 'D+D×Sex' – females: r = 0.54, p = 0.003, males: r = 0.52, p = 0.005).

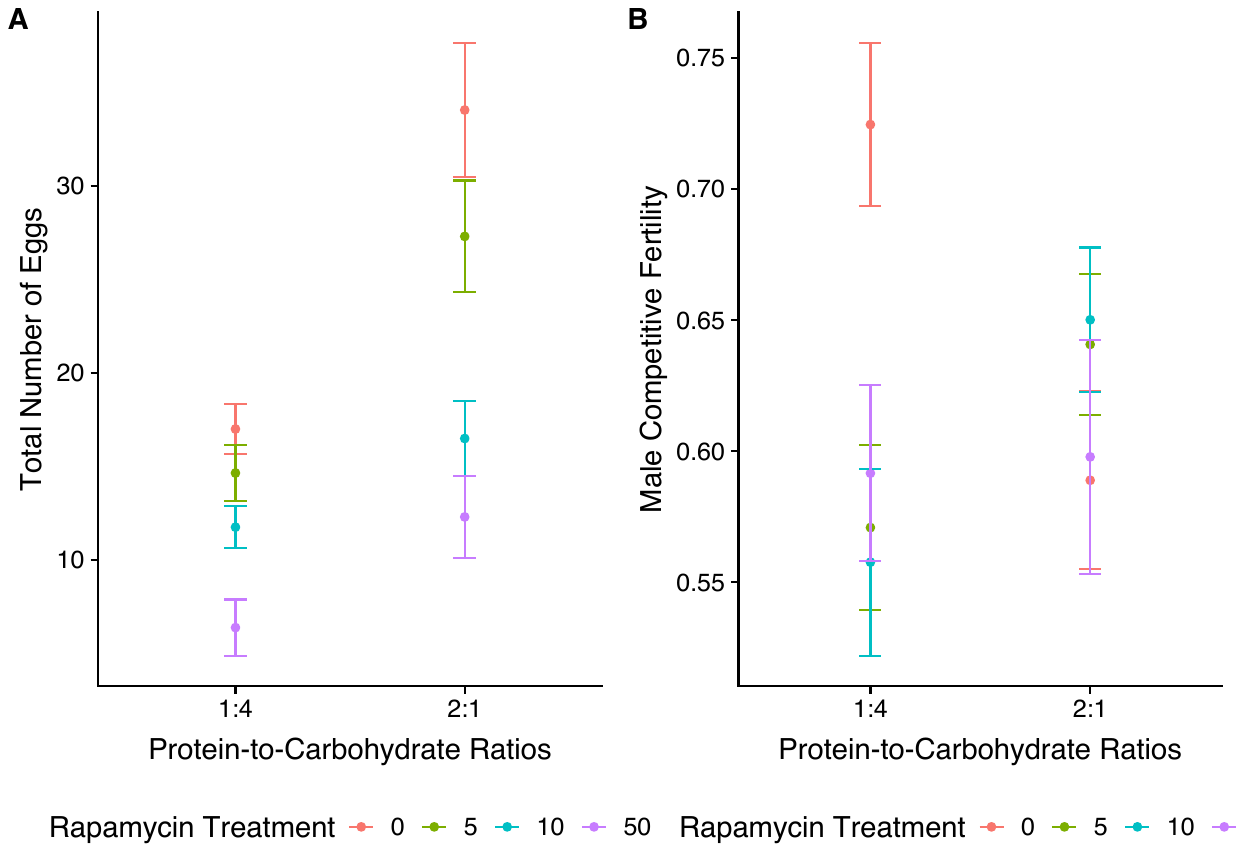

Female Fecundity

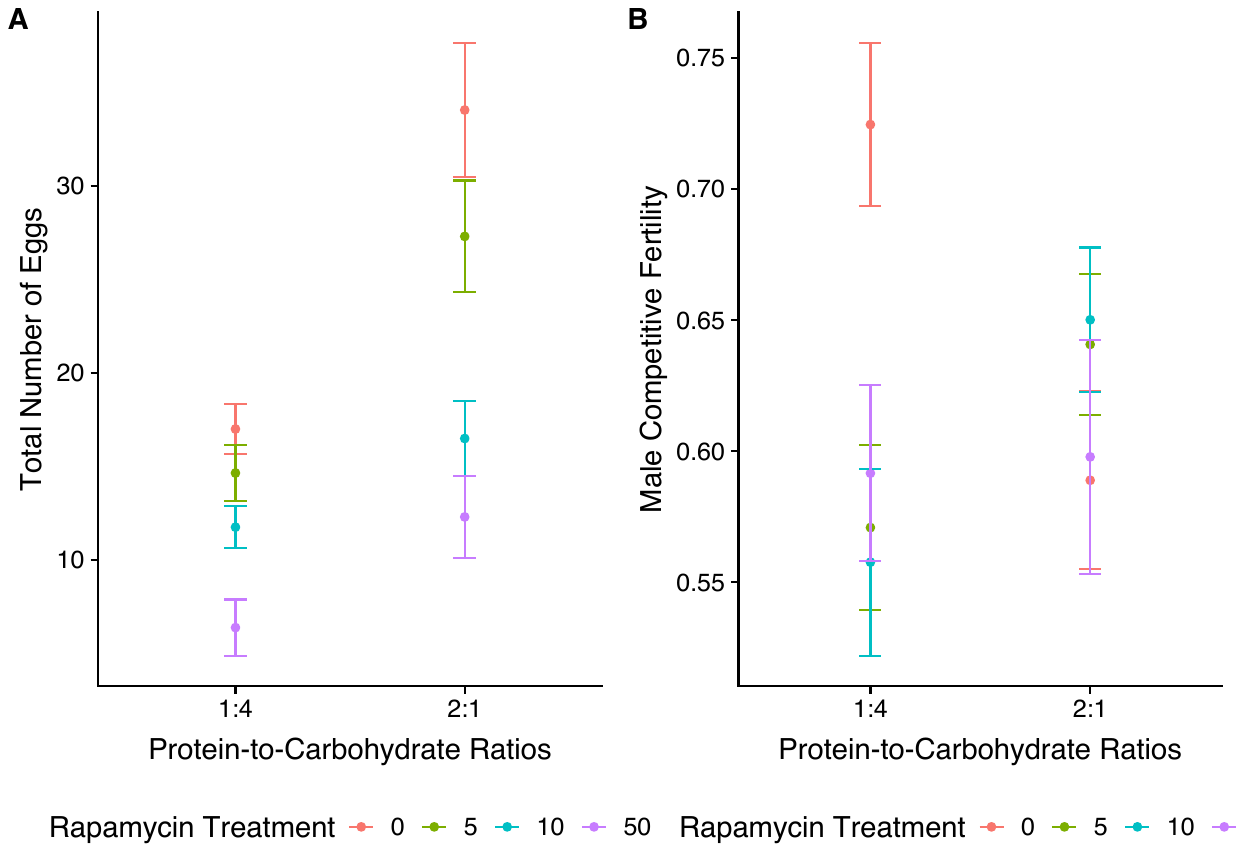

**Figure S7:** Sex-specific fitness measured across both diets and 4 rapamycin treatments. Panel (A) shows data for female fecundity, panel (B) data for male competitive fertility.
